# Resting high frequency heart rate variability is not associated with the recognition of emotional facial expressions in healthy human adults

**DOI:** 10.1101/077784

**Authors:** Brice Beffara, Nicolas Vermeulen, Martial Mermillod

**Affiliations:** Univ. Grenoble Alpes, LPNC, F-38040, Grenoble, France; CNRS, LPNC UMR 5105, F-38040, Grenoble, France; IPSY, Universitè Catholique de Louvain, Louvain-la-Neuve, Belgium; Fund for Scientific Research (FRS-FNRS), Brussels, Belgium

**Keywords:** HF-HRV, autonomic flexibility, emotion identification, dynamic EFEs, Polyvagal theory, Neurovisceral integration model

## Abstract

This study explores whether the myelinated vagal connection between the heart and the brain is involved in emotion recognition. The Polyvagal theory postulates that the activity of the myelinated vagus nerve underlies socio-emotional skills. It has been proposed that the perception of emotions could be one of this skills dependent on heart-brain interactions. However, this assumption was differently supported by diverging results suggesting that it could be related to confounded factors. In the current study, we recorded the resting state vagal activity (reflected by High Frequency Heart Rate Variability, HF-HRV) of 77 (68 suitable for analysis) healthy human adults and measured their ability to identify dynamic emotional facial expressions. Results show that HF-HRV is not related to the recognition of emotional facial expressions in healthy human adults. We discuss this result in the frameworks of the polyvagal theory and the neurovisceral integration model.

## Introduction

The behavior of an animal is said social when involved in interactions with other animals (Ward & Webster, 2016). These interactions imply an exchange of information, signals, between at least two animals. In humans, the face is an efficient communication channel, rapidly providing a high quantity of information. Facial expressions thus play an important role in the transmission of emotional information during social interactions. The result of the communication is the combination of transmission from the sender and decoding from the receiver (Jack & Schyns, 2015). As a consequence, the quality of the interaction depends on the ability to both produce and identify facial expressions. Emotions are therefore a core feature of social bonding (Spoor & Kelly, 2004). Health of individuals and groups depend on the quality of social bonds in many animals (Boyer, Firat, & Leeuwen, 2015; S. L. Brown & Brown, 2015; Neuberg, Kenrick, & Schaller, 2011), especially in highly social species such as humans (Singer & Klimecki, 2014).

The recognition of emotional signals produced by others is not independent from its production by oneself (Niedenthal, 2007). The muscles of the face involved in the production of a facial expressions are also activated during the perception of the same facial expressions (Dimberg, Thunberg, & Elmehed, 2000). In other terms, the facial mimicry of the perceived emotional facial expression (EFE) triggers its sensorimotor simulation in the brain, which improves the recognition abilities (Wood, Rychlowska, Korb, & Niedenthal, 2016). Beyond that, the emotion can be seen as the body -including brain-dynamic itself (Gallese & Caruana, 2016) which helps to understand why behavioral simulation is necessary to understand the emotion.

The interplay between emotion production, emotion perception, social communication and body dynamics has been summarized in the framework of the polyvagal theory (Porges, 2007). In a phylogenetic perspective, the polyvagal theory describes how the interaction between the central and the autonomic nervous systems underlie social behaviors. Heart brain interactions are the core feature of the theory because they shape the adaptation of an organism to environmental variations. Indeed, social interactions precisely generate a large amount of variability in the environment (Taborsky & Oliveira, 2012). Three major phylogenetic stages are identified in the polyvagal theory and are all associated with a specific physiological functioning. The most primitive stage is supposed common to almost all the vertebrates. The behavioral function associated is immobilization and is underpinned by the unmyelinated branch of the vagus nerve connecting the heart and the brain. This function is a defense mechanism allowing to cope with highly dangerous events. The fight/flight response to danger is assumed to have emerged during a second and more recent stage and is dependent on the sympathetic-adrenal system. Finally, the third and last stage is proposed to characterize most of the mammals. The major physiological component of this stage is the myelinated branch of the vagus nerve which underlies self-soothing and prosocial/affiliative behaviors.

The myelinated vagus nerve quickly conducts information between heart and brain resulting in modifications of heart rate and heart contraction (Coote, 2013). The vagus nerve and the heart are connected at the level of the sinus node via acetylcholine. The sinus node contains high quantity of acetylcholinesterase, the acetylcholine is rapidly hydrolyzed, and the delay of vagal inputs are short (Task Force of the European Society of Cardiology the North American Society of Pacing Electrophysiology (1996);Thayer2012a). Secondly, myelinated axonal conduction speed is high resulting in a quick reaction of the heart to the stimulation and to the stop of the stimulation (T. W. Ford & McWilliam, 1986; Jones, Wang, & Jordan, 1995; D. Jordan, 2005). High speed communication between the heart and the brain generates important variability in heart rate. This physiological variability allowed by vagal activity contributes to optimal regulation of the metabolism as a function of environmental changes and internal needs (Porges, 1997; Thayer & Sternberg, 2006).

Axons of the myelinated vagus nerve originates from preganglionic cardiac vagal neurons situated in the nucleus ambiguus (Porges, 1997). The nucleus ambiguus is a group of motor neurons from which the myelinated branch but also several sensory and motor fibers including the facial and trigeminal nerves emerge (Porges, 1998). The nucleus ambiguus has bidirectional connections with cortical (prefrontal, cingulate and insular) and sub-cortical (amygdala, hypothalamus) areas (Thayer & Lane, 2009). Theses regions play an important role in social cognition (Amodio & Frith, 2006) and emotional processing (Lane et al., 2009). This implies that social communication, cardiac vagal control and facial muscular control share common structural pathways. Constantly receiving updated information from external and internal changes, the nucleus ambiguus is the place of rapid central-periphery integration and reactivity toward emotional challenges (Coote, 2013; Porges, 1995). Afferent inputs to the facial motor nucleus are found (inter alia) in the nucleus ambiguus. The distribution of motoneurons supplying fast muscle contraction might underlie the complexity and mobility of facial expressions (Sherwood, 2005). The panel of available facial expressions could be the result of dynamic connections between cortical control, brainstem nuclei sensorimotor integration/inhibition and facial muscles activity (Porges, 2001) and may foster the ability to engage and regulate diversified social interactions (Sherwood, 2005). It is to notice that this proposition made by the polyvagal theory (Porges, 2001) seems plausible but has not been developed or tested specifically at an anatomo-functional level. Indeed, an important gap remains between the functions of neural connections and social skills. Evidence toward this hypothesis is mitigated so far (Sherwood, 2005) and needs to be tested further both at anatomical, physiological and behavioral levels.

Taken together, anatomo-functional characteristics of heart-brain-face interactions allow to predict that myelinated vagus nerve activity should be associated with the ability to process emotional facial signals involved in social communication (Porges, 2003). However, even if the literature cited above strongly corroborate the hypothesis formulated by Porges (1995), measures of vagal activity and emotion signal perception have not been recorded together until Bal et al. (2010) in healthy children and autistic children and Quintana, Guastella, Outhred, Hickie, & Kemp (2012) in healthy human adults. They monitored the myelinated vagal heart-brain communication via the spectral analysis of heart rate variability which is a popular and reliable non-invasive tool reflecting the autonomic nervous system activity (Heathers, 2014; Task Force of the European Society of Cardiology the North American Society of Pacing Electrophysiology, 1996). Specifically, they extracted high frequency range of heart rate variability (HF-HRV) which provides a rigorous assessment of the myelinated heart-brain connection activity [Akselrod et al. (1981); Gary G Berntson et al. (1997); G. G. Berntson, Cacioppo, & Quigley (1993); Gary G. Berntson, Norman, Hawkley, & Cacioppo (2008); Cacioppo et al. (1994); M V Kamath & Fallen (1993); M. V. Kamath, Upton, Talalla, & Fallen (1992); M V Kamath, Upton, Talalla, & Fallen (1992)).

Bal et al. (2010) evaluated facial emotion recognition with videos of dynamic EFEs (Dynamic Affect Recognition Evaluation, DARE) on the six basic emotions (sadness, fear, surprise, disgust, anger, and happiness, Porges, Cohn, Bal, & Lamb (2007)). Videos displayed emotions going from neutral expression to apex through morphing. Quintana et al. (2012) evaluated facial emotion recognition by the Reading Mind in the Eyes Test (RMET, (Baron-Cohen, Jolliffe, Mortimore, & Robertson, 1997; Baron-Cohen, Wheelwright, Hill,Raste, & Plumb, 2001)). The RMET is composed of photographs displaying the eye-region of the facial expression of actors/actresses. The facial expressions corresponds to a feeling, a thinking or mental state. The photograph is displayed along with 4 labels describing possible mental states, among which only one actually corresponds to the picture. The task of the participants is therefore to “read in the mind” in order to identify the correct mental state. The work of Quintana et al. (2012) is important because no data could bring evidence in favor of an association between vagal activity and the perception of social cues in healthy humans within the polyvagal framework (Porges, 1997) until them. The main result of their study is that HF-HRV is associated with better scores at the RMET, with items recoded such as correct answers weight much for difficult trials versus easy ones. The authors conclude that higher levels of resting state HF-HRV are associated with better emotion recognition skills. Conversely, Bal et al. (2010) did not find any association between HF-HRV and emotion identification in healthy participants but only in children with autism spectrum disorders. Besides this results is observed only for response latency but not for accuracy). Fear, happiness and sadness were faster identified by higher resting-state HF-HRV participants.

On one side, Quintana et al. (2012) found that healthy adults were better at identifying mental states (when weighting for difficulty) and therefore proposed that HF-HRV is linked with emotion recognition. One the other side, Bal et al. (2010) did not find any association between HF-HRV and emotion identification (in healthy children). From here, 2 explanations can emerge: i) The conceptual overlap between emotion recognition and mind reading found in Quintana et al. (2012) matters, and HF-HRV is associated with mental state reading bu not with emotion identification *per se*, iii) Considering healthy human participants, HF-HRV is associated with emotion recognition only in adults.

A study mixing the designs of Bal et al. (2010) and Quintana et al. (2012) can help to disentangle between these hypotheses. We report the results obtained after a protocol where resting state HF-HRV is measured in healthy adults. The emotion identification task is similar to the Dynamic Affect Recognition Evaluation software (DARE, Porges et al. (2007)) used in Bal et al. (2010) (including anger, disgust, fear, joy, sadness and surprise) but included three more EFEs (contempt, embarrassment, and pride). All EFEs movies were from the Amsterdam Dynamic Facial Expression Set (ADFES, Schalk, Hawk, Fischer, & Doosje (2011)), a more recent database with color stimuli. As a consequence, emotion identification is based on a recent database with dynamic EFEs used in Bal et al. (2010) (anger, disgust, fear, joy, sadness and surprise) and 3 more (contempt, embarrassment, and pride) in order to increase complexity. Even if this perspective is strongly challenged (Jack, Sun, Delis, Garrod, & Schyns, 2016), some authors suggest that the emotions used by Bal et al. (2010) are more basic and easier to identify compared to emotions more complex emotions such as contempt, embarrassment, and pride (Baron-Cohen, Golan, & Ashwin, 2009). Contempt, embarrassment and pride are considered as “self-conscious” emotions but present typical morphological configurations at the level of the whole face (Schalk et al., 2011). Indeed, they involve facial muscular patterns or even slight movement of the head (Tracy & Robins, 2008; Tracy, Robins, & Schriber, 2009) and these patterns are to be decoded in order to identify the emotion. Albeit more complex than basic emotions, they differ from pure mental states because not concentrated on the eyes area.

As the distinction between basic and complex emotions fits the difference between our set of EFEs and the set used by Bal et al. (2010), it is relevant to rely on it as a factor of difficulty in EFEs recognition. Our design allows to assess if HF-HRV is associated with emotion recognition on a new set of dynamic whole EFEs. If HF-HRV is associated with emotion recognition in these conditions, this suggests that the task used by Bal et al. (2010) was not complex enough to establish the correlation and that HF-HRV is not discriminant for the recognition of “basic” emotions. On the contrary, if HF-HRV is not associated with emotion recognition, this would suggest that the results of Quintana et al. (2012) does not apply to emotion perception *per se* but rather to different “non-emotional” mechanisms involved in social signals reading (R. L. C. Mitchell & Phillips, 2015).

## Methods

In the “Methods” and “Data analysis” sections, we report how we determined our sample size, all data exclusions, all manipulations, and all measures in the study (Simmons, Nelson, & Simonsohn, 2012).

### Sample

Initial sample was composed of 77 young healthy human adults. Participants were recruited via advertisements (mailing list and poster). All participants were psychology students of University Grenoble-Alpes. Participants were French or perfectly bilingual in French. They provided written informed consent before the participation. The study was part of a global project reviewed and approved by the University human ethics committee from Grenoble, France (Grenoble ethics committee notice number 2014-05-13-49 and 2014-05-13-48). To be eligible, participants had to be aged between 18 and 60 years, with a normal or normal-to-corrected vision, explicitly reported an absence of psychiatric, neurologic, hormonal, or cardio-vascular disease, and with no medical treatment (with the exception of contraception). Smoking, energizing drinks (e.g. coffee, tea, etc…) and psychotropic substances (e.g. alcohol, cannabis, etc…) were prohibited to each participant the day of the experiment. They had also to avoid eating or drinking (water was allowed) the 2 hours preceding the experiment in order to limit the influence of digestion on autonomic functioning (Short term HRV measurement can be biased by the digestion of food since viscera are innervated by the autonomic nervous system (Heathers, 2014; Iorfino, Alvares, Guastella, & Quintana, 2016; Quintana & Heathers, 2014)) but they had to eat in the morning (more than 2 hours before the experiment) in order to avoid fasting states. The participants received experimental credits in return of their participation.

### Sample size

We planned between 75 and 80 participants to take part in the study. Anticipating possible exclusions due to technical problems, we determined our sample size expecting at least 65 participants suitable for final analysis. This sample size was set on the basis of Quintana et al. (2012). Their sample size of 65 was adequate to observe an association between HF-HRV and the RMET score, with an effect size of *R*^2^~.07.

### Procedure

The experiment took place in a quiet and dimmed room. All participants were tested between 0900 h and 1300 h. After a global description of the experiment, participants were asked to empty their bladder before starting the experiment. After that, they were taught how to install the Bioharness^®^ heart rate monitor. They were left in autonomy in an isolated room for the installation of the heart rate monitor. Then, they seated in a chair, the experimenter checked the signal and the experiment started. The instructions were to relax, breathe naturally and spontaneously. During 5 minutes, the participant watched short neutral samples of films selected and evaluated by Hewig et al. (2005) (“Hannah and her Sisters” and “All the President’s Men”) and Schaefer, Nils, Sanchez, & Philippot (2010) (“Blue [1]”, “Blue [2]”, “Blue [3]” and “The lover”). Videos were displayed without audio. These 5 first minutes aimed to allow the participant to shift in a calm state. ECG data for HRV baseline computation was recorded during the 5 following minutes while participants listened to the first 5 minutes of a neutral audio documentary designed for laboratory studies (Bertels, Deliens, Peigneux, & Destrebecqz, 2014). Neutral videos and audio documentary were used in order to standardize ECG recordings (Piferi, Kline, Younger, & Lawler, 2000). ECG data was recorded during spontaneous breathing (Denver, Reed, & Porges, 2007; Kobayashi, 2009; Kowalewski & Urban, 2004; Larsen, Tzeng, Sin, & Galletly, 2010; Muhtadie, Koslov, Akinola, & Mendes, 2015; Pinna et al., 2007). After this first phase, the emotion identification task session started for about 15 minutes (see description below). When this step ended, the participant completed computerized control surveys. The experimenter stayed out the room during the experiment but was available for eventual questions between the different steps of the experiment.

### Emotion identification task

The emotion identification task followed the design used by Bal et al. (2010) and proposed by Porges et al. (2007). Participants were presented with short video clips displaying dynamic standardized EFEs produced by humans adults. All the stimuli came from the ADFES (Schalk et al., 2011). Nine EFEs (Figure 1) of ten North-European models (5 males and 5 females: “F01”, “F02”, “F03”, “F04”, “F05”, “M02”, “M03”, “M04”, “M11”,and “M12”) were presented in a random design. Video clips displayed the face of the model going from a neutral expression to the apex of the EFE. Video clips duration ranged from 6 to 6.5 seconds, including a neutral face for 0.5 seconds, followed by the onset of the EFE, and then the face held at apex for 5 seconds (Figure 2). In phase 1 of each trial, participants used the numeric pad of the computer keyboard to identify EFEs. The were asked to push the “0” key as soon as they could identify what emotion was expressed in the video video clip. Synchronous with the “0” key press, phase 2 started as the the video clip stopped and a new screen appeared with each of the nine emotion labels matched with one the nine other number keys of the numeric pad (1-2-3-4-5-6-7-8-9). The same matching – randomly determined before the launch of the experiment – was used for all trials and for all participants (pride = 1, sadness = 2, surprise = 3, embarrassment = 4, fear = 5, joy = 6, anger = 7, contempt = 8, disgust = 9). The participant was asked to identify which of the nine emotion labels corresponded to the EFE displayed in the video clip. There was no time limit nor time pressure or measure for phase 2 responses. The latency to recognize emotions was measured as the response time in phase 1. Emotion recognition accuracy was measured by the responses provided during phase 2. Before data recording, participants performed nine training trials on the nine EFEs of another North-European model (“M08”) in order to familiarize with the task. The experimenter stayed in the experimental room during this step in order to check if the participant understood the instructional set and possibly help the participant in case of questions and/or difficulties.

**Figure 1.**
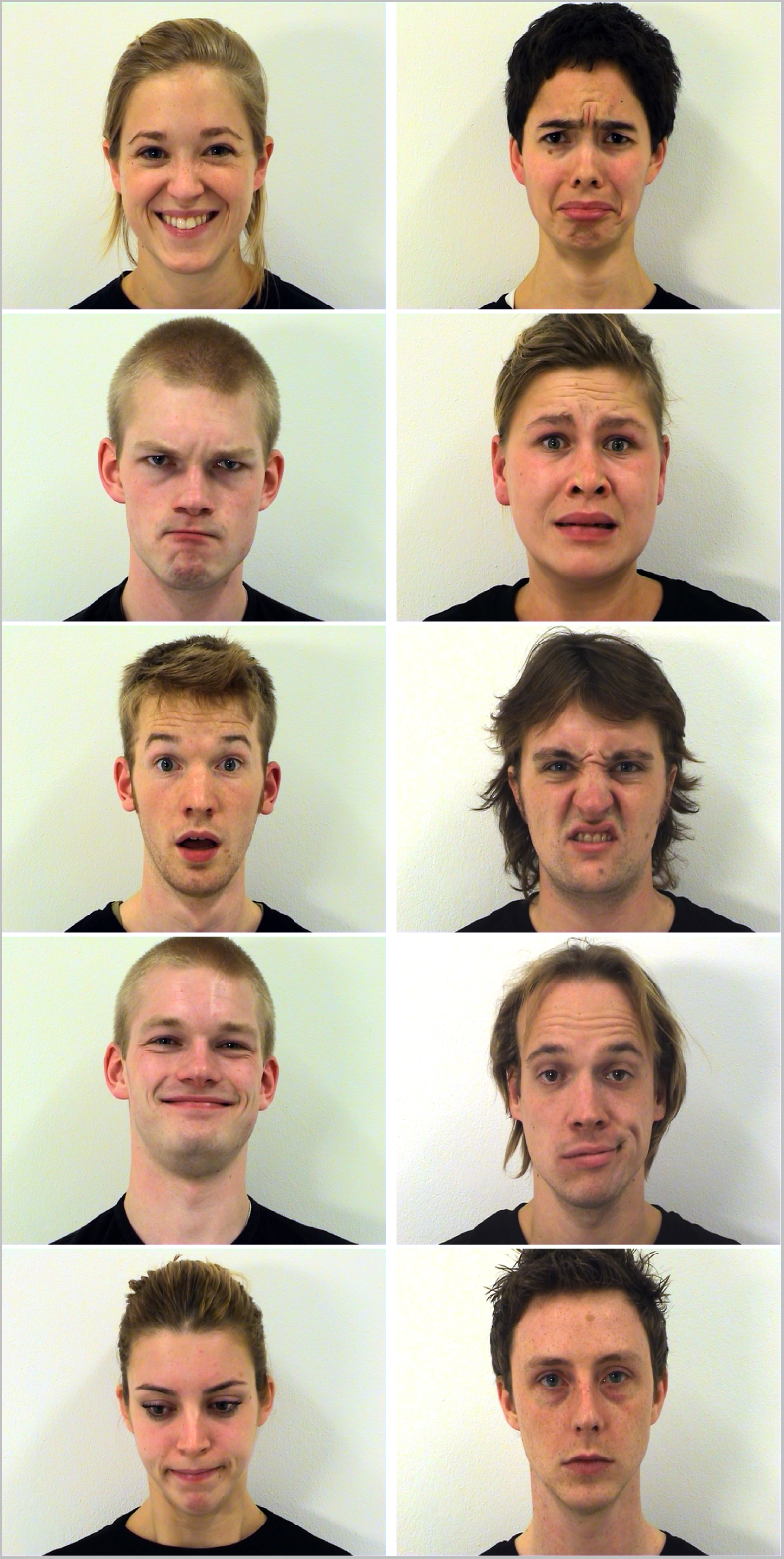
Examples of emotional facial expressions and of a neutral facial expression. From left to right and top to bottom: Joy (F03), Sadness (F04), Anger (M03), Fear (F05), Surprise (M02), Disgust (M04), Pride (M03), Contempt (M11), Embarras (F01), and Neutral (M12). All stimuli are from the ADFES (van der Schalk, Hawk, Fischer, & Doosje, 2011).

**Figure 2.**
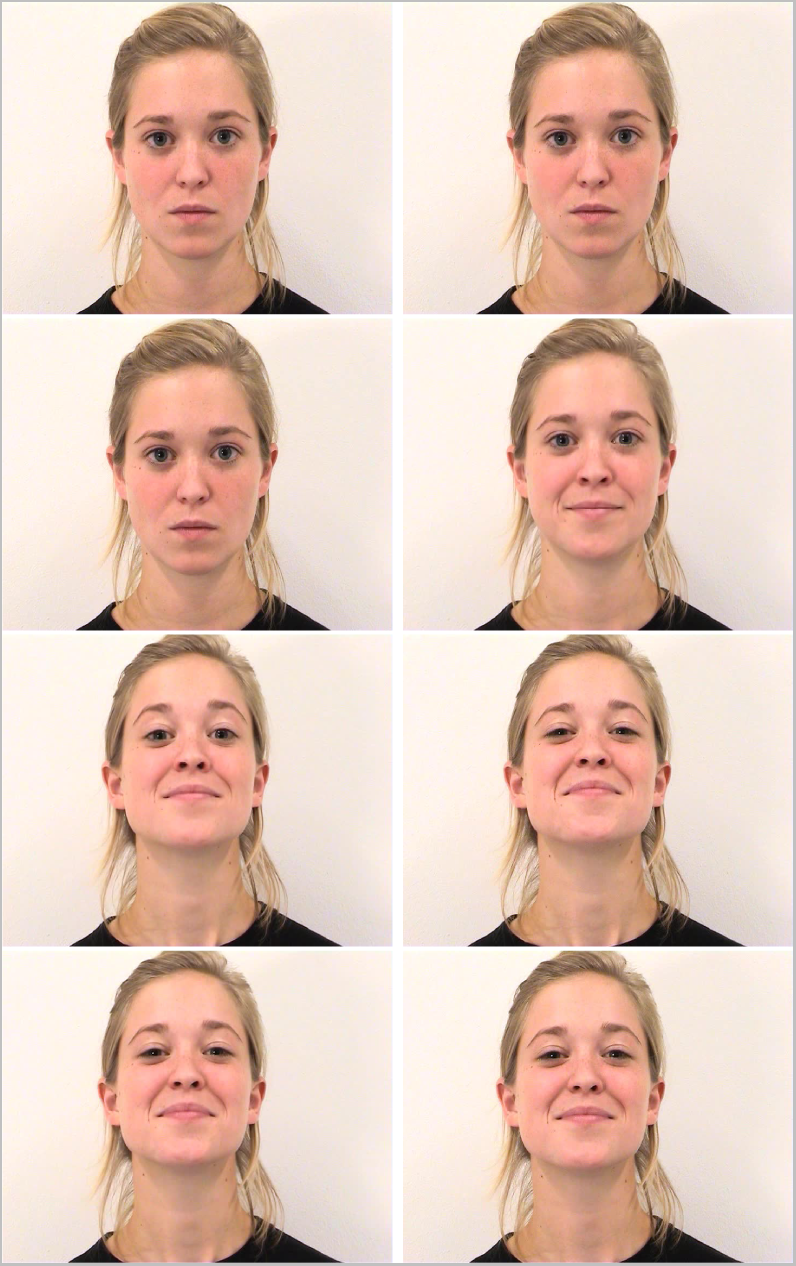
Time course of the EFE of pride (F03). From left to right and top to bottom, t = 0, 0.5, 1, 2, 3, 4, 5, 6 seconds. Images are extracted from the original video set of the ADFES (van der Schalk, Hawk, Fischer, & Doosje, 2011)

Response times were used as a measure of quantity of evidence needed in order to detect the emotion (Bal et al., 2010). In other words, the hypothesis behind is that more efficient processing of facial emotional signals should allow to detect more subtle muscle movements of the face and therefore identify the emotion faster. Obviously, this method could also be influenced by different strategies of response but accuracy scores are available in order to assess the success of the recognition. As a consequence, performances in emotion recognition can be evaluated by both measures separately. The presence of 9 instead of 6 emotions (compared to Bal et al. (2010)) allows to increase the difficulty of the task and therefore induce variability in our data. Therefore, this design is closer to the design proposed by Quintana et al. (2012) with a large number of different emotions to categorize.

### Physiological measurement

The electrocardiogram (ECG) data was recorded with a Zephyr Bioharness^TM^ 3.0 (Zephyr, 2014). The Bioharness^TM^ is a class II medical device presenting a very good precision of measurement for ECG recording in low physical activity conditions (Johnstone, Ford, Hughes, Watson, & Garrett, 2012a, 2012b; Johnstone et al., 2012). It has been used for ECG measurements in both healthy and clinical populations, presenting a very high-to-perfect correlation with classical hospital or laboratory devices (Brooks et al., 2013; Yoon, Shah, Arnoudse, & De La Garza, 2014). The Bioharness^TM^ both provides comfort for the participant and allow reliable HRV extraction for the researcher (Lumma, Kok, & Singer, 2015). The chest strap’s sensor measures electrical activity corresponding to the classical V4 lead measurement (5th intercostal space at the midclavicular line) through conductive Lycra fabric. A single-ended ECG circuit detects QRS complexes and incorporates electrostatic discharge protection, both active and passive filtering and an analog-to-digital converter. Interbeat intervals are derived by Proprietary digital filtering and signal processed with a microcontroller circuit. The ECG sensor sampling frequency is 250 Hz and the resolution 0.13405 mV., ranging from 0 to 0.05 V (Villarejo, Zapirain, & Zorrilla, 2013). After a slight moistening of the 2 ECG sensors, the chest-strap was positioned directly on the skin, at the level of the inframammary fold, under the lower border of the pectoralis major muscle. The recording module communicated with an Android^®^ OS smartphone by Bluetooth^®^. The application used to acquire the signal emitted by the Bioharness^TM^ was developed, tested, and validated by Cânovas, Domingues, & Sanches (2011). The Android^®^ OS device used to record the signal was an LG-P990 smartphone (Android^®^ version 4.1.2.).

### Control for confounding factors

To control for confounding variables likely to be linked to HRV, participants completed questionnaires detailing life habits, demographic data and emotional traits (Quintana et al., 2012). Physical activity was assessed with the International Physical Activity Questionnaire (IPAQ,Craig et al. (2003)), composed of 9 items that calculate an index reflecting the energy cost of physical activities (Metabolic Equivalent Task score, MET). The IPAQ has been validated in French (Briancon et al., 2010; Hagströmer, Oja, & Sjöström, 2006) and widely used in French surveys (Salanave et al., 2012). Participants also completed the Depression Anxiety and Stress scales (DASS-21;(P. F. Lovibond & Lovibond, 1995)). The DASS-21 is a 21-item questionnaire, validated in French (Ramasawmy & Gilles, 2012), and composed of three subscales evaluating depression, anxiety and stress traits. We also recorded the size, weight, age and sex of the participants and their daily cigarette consumption. Participants answered final surveys on a DELL latitude E6500 laptop. Surveys were built and displayed with E-prime software (E-prime 2.0.10.242 pro).

### Physiological signal processing

R-R interval data was extracted from the Android^®^ device and imported into RHRV for Ubuntu (Rodríguez-Liñares et al., 2011). Signal was visually inspected for artifact (Prinsloo et al., 2011; Quintana et al., 2012; Wells, Outhred, Heathers, Quintana, & Kemp, 2012). Ectopic beats were discarded (Kemper, Hamilton, & Atkinson, 2007) for participants presenting a corrupted RR interval series (Beats per minute (bpms) shorter/longer than 25/180 and/or bigger/smaller than 13% compared to the 50 last bpms). RR series were interpolated by piecewise cubic spline to obtain equal sampling intervals and regular spectrum estimations. A sampling rate of 4 Hz was used. We then extracted the frequency component of HRV from RR interval data. The LF (0.04-0.15 Hz) and HF (0.15-0.4 Hz) components were extracted using an east asymmetric Daubechies wavelets with a length of 8 samples. Maximum error allowed was set as 0.01 (García, Otero, Vila, & Márquez, 2013).

#### Model comparison

Model selection was completed using *AIC*_*c*_ (corrected Akaike information criterion) and Evidence Ratios -*ER*_*i*_- (K. P. Burnham & Anderson, 2004; Kenneth P. Burnham, Anderson, & Huyvaert, 2011; Hegyi & Garamszegi, 2011; Symonds & Moussalli, 2011). *AIC*_*c*_ provides a relative measure of goodness-of-fit but also of parsimony by sanctioning models for their numbers of parameters. *AIC*_*c*_ is more severe on this last point than 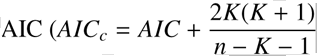 where *K* is the number of parameters and *n* the sample size.). We computed the difference between best (lower) and other *AIC*_*c*_ *s* with Δ_*AIC*_*c*__ = *AIC*_*c*_*i*__ − *AIC*_*c*_*min*__. The weight of a model is then expressed as 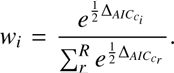. From there,we can compute the Evidence Ratio: 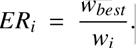. Even if quantitative information about evidence is more precise, we also based our decision on Kass & Raftery (1995) and Snipes & Taylor (2014), i.e. minimal (*ER*_*i*_ < 3.2), substantial (3.2 < *ER*_*i*_ < 10), strong (10 < *ER*_*i*_ < 100) and decisive (100 < *ER_i_*) evidence. If the model with the lower *AIC*_*c*_ included more parameters than others, we considered it as relevant if the evidence was at least substantial. If the model with the lower *AIC*_*c*_ included less parameters than others, we chose it even if evidence was minimal.

## Results

Correlations between control variables and variables of interest are displayed in figures 3 and 4. Because weight was associated with HF-HRV, we adjusted HF-HRV for it by extracting the standardized residuals of the regression with weight as the independent variable and HF-HRV as the dependent variable (Quintana et al., 2012). HRV as an independent variable in the following analysis is therefore HF-HRV (normalized units) adjusted for weight.

**Figure 3.**
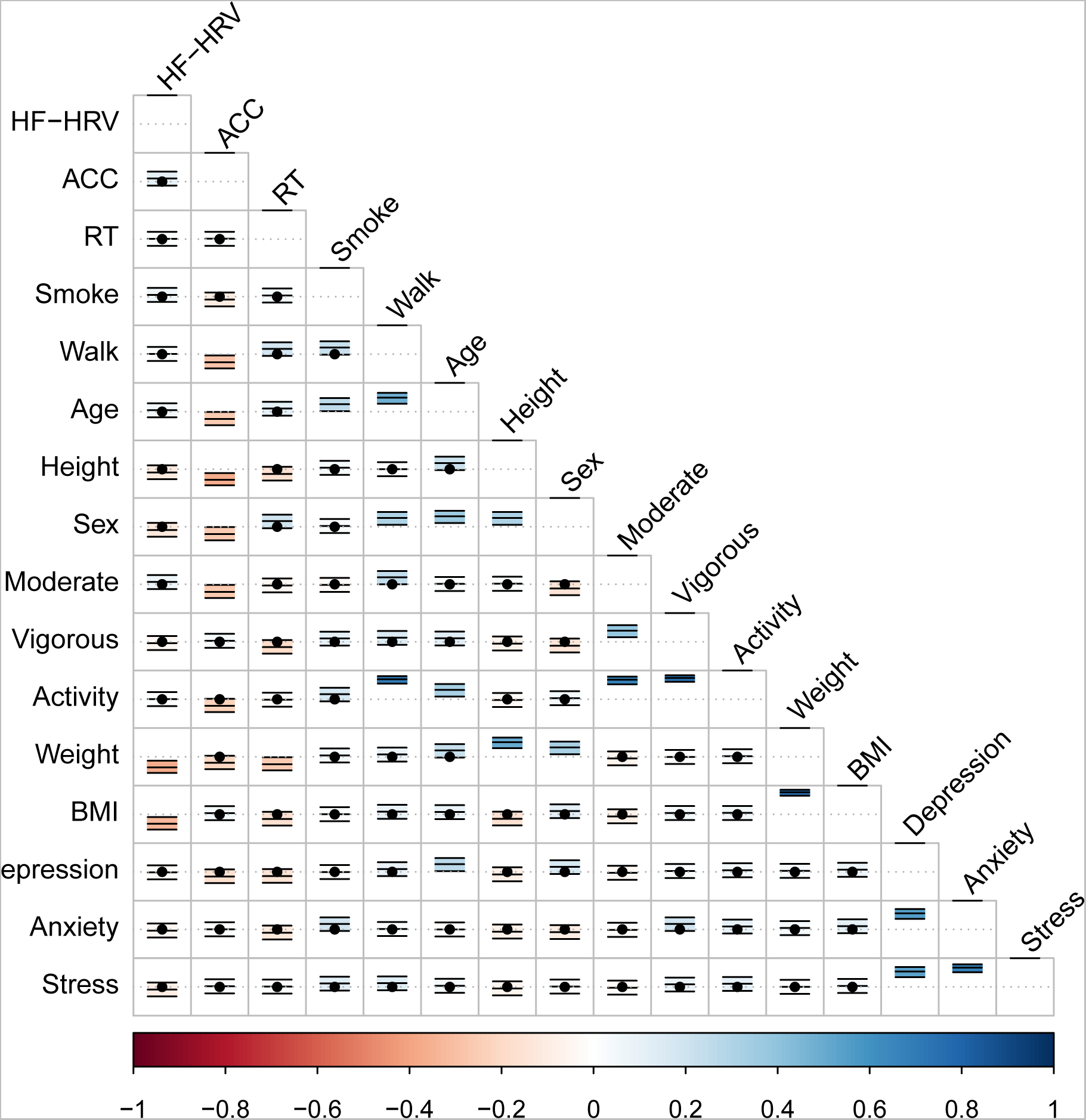
Correlation confidence intervals between recorded variables. Confidence regions represent 95% CIs and are marked with a black dot when including 0.

**Figure 4.**
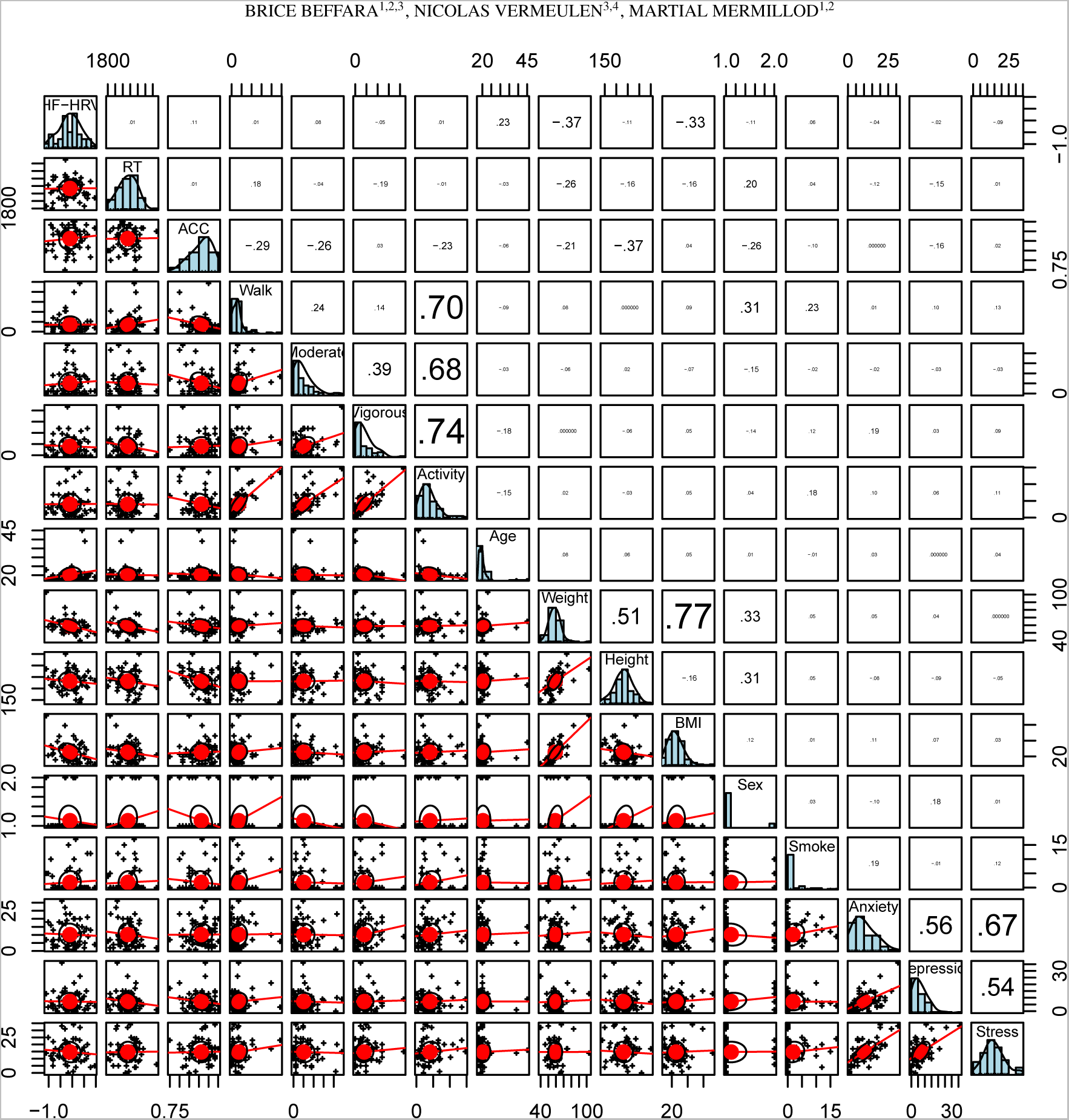
Scatter plots, distributions, and Pearson correlation coefficients between recorded variables. R values’ font sizes are proportional to the strength of the correlation

In a second step, we selected the relevant random factors to include in our models. Whether for response times or accuracy, participants and items (i.e. the model (actor) performing the EFE) were appropriate as random factors. Indeed models including participants and items showed the lowest (best) AICc with *ER*_*i*_ = 0.936/0.064 = 14.62 (strong evidence) for response times (Table 1) and *ER*_*i*_ = 0.967/0.033 = 29.30 (strong evidence) for accuracy (Table 2).

**Table 1.**
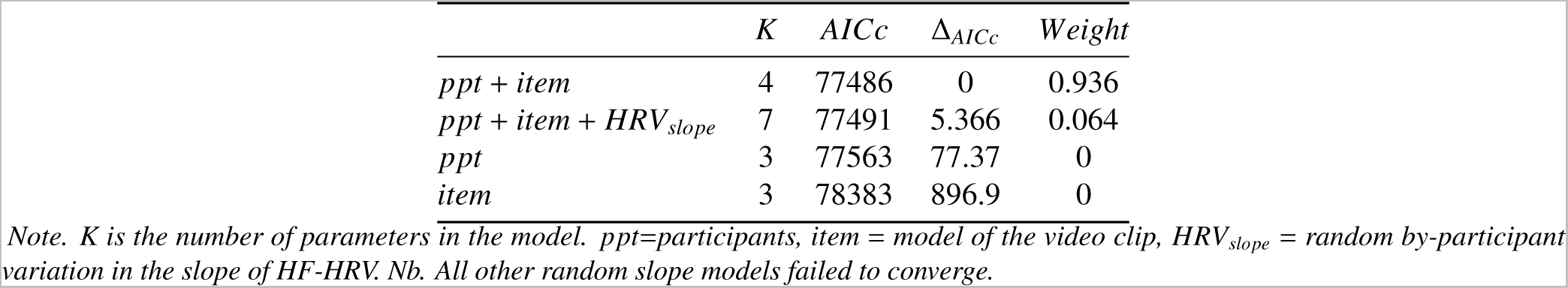
Comparison of random effects in models for response times, ordered by AICc relative to the model with the lowest (best) AICc.

| | *K* | *AICc* | $\Delta_{AICc}$ | *Weight* |
| --- | --- | --- | --- | --- |
| *ppt + item* | 4 | 77486 | 0 | 0.936 |
| *ppt + item + HRV<sub>slope</sub>* | 7 | 77491 | 5.366 | 0.064 |
| *ppt* | 3 | 77563 | 77.37 | 0 |
| *item* | 3 | 78383 | 896.9 | 0 |

Note. *K* is the number of parameters in the model. *ppt*=participants, *item* = model of the video clip, *HRV<sub>slope</sub>* = random by-participant variation in the slope of HF-HRV. Nb. All other random slope models failed to converge.

**Table 2.** Comparison of random effects in models for accuracy, ordered by AICc relative to the model with the lowest (best) AICc.

| | *K* | *AICc* | $\Delta_{AICc}$ | *Weight* |
| --- | --- | --- | --- | --- |
| *ppt + item* | 3 | 4463 | 0 | 0.967 |
| *ppt + item + HRV<sub>slope</sub>* | 6 | 4469 | 6.739 | 0.033 |
| *ppt* | 2 | 4494 | 31.66 | 0 |
| *item* | 2 | 4516 | 53.21 | 0 |

Note. *K* is the number of parameters in the model. *ppt*=participants, *item* = model of the video clip, *HRV<sub>slope</sub>* = random by-participant variation in the slope of HF-HRV. Nb. All other random slope models failed to converge.

**Table 3.**
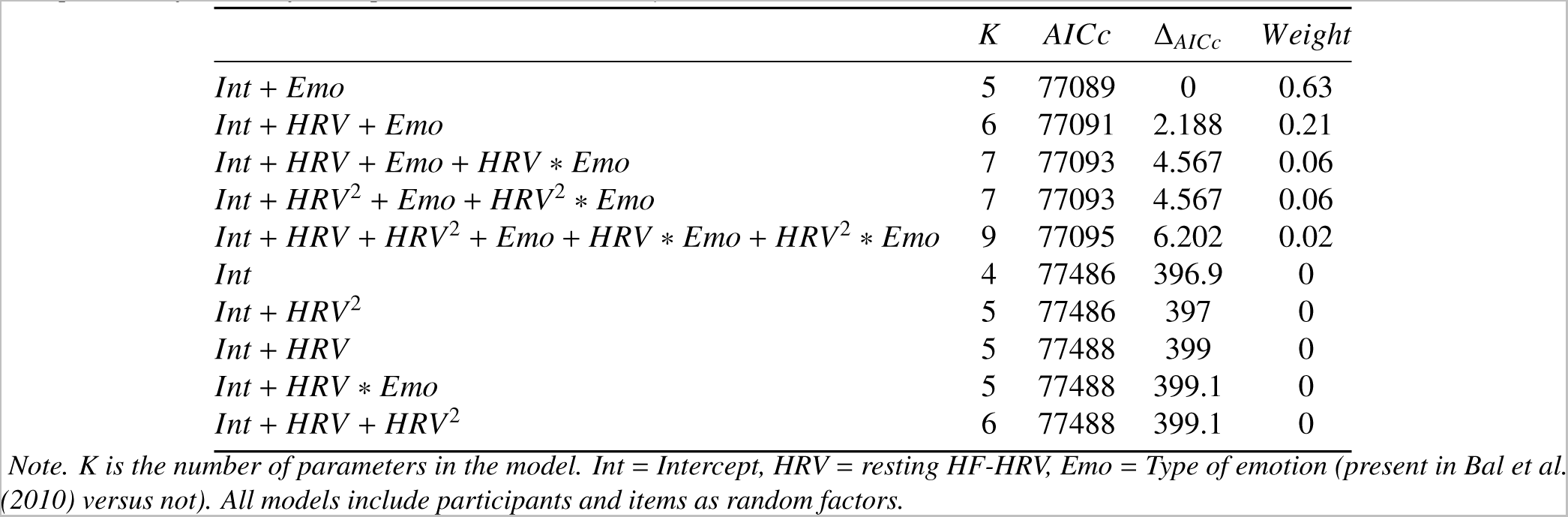
Comparison of models for response times, ordered by AICc relative to the model with the lowest (best) AICc.

| | *K* | *AICc* | $\Delta_{AICc}$ | *Weight* |
| --- | --- | --- | --- | --- |
| *Int + Emo* | 5 | 77089 | 0 | 0.63 |
| *Int + HRV + Emo* | 6 | 77091 | 2.188 | 0.21 |
| *Int + HRV + Emo + HRV \* Emo* | 7 | 77093 | 4.567 | 0.06 |
| *Int + HRV<sup>2</sup> + Emo + HRV<sup>2</sup> \* Emo* | 7 | 77093 | 4.567 | 0.06 |
| *Int + HRV + HRV<sup>2</sup> + Emo + HRV \* Emo + HRV<sup>2</sup> \* Emo* | 9 | 77095 | 6.202 | 0.02 |
| *Int* | 4 | 77486 | 396.9 | 0 |
| *Int + HRV<sup>2</sup>* | 5 | 77486 | 397 | 0 |
| *Int + HRV* | 5 | 77488 | 399 | 0 |
| *Int + HRV \* Emo* | 5 | 77488 | 399.1 | 0 |
| *Int + HRV + HRV<sup>2</sup>* | 6 | 77488 | 399.1 | 0 |

Note. *K* is the number of parameters in the model. *Int* = Intercept, *HRV* = resting HF-HRV, *Emo* = Type of emotion (present in Bal et al. (2010) versus not). All models include participants and items as random factors.

**Table 4.** Comparison of models for accuracy, ordered by AICc relative to the model with the lowest (best) AICc.

| | *K* | *AICc* | $\Delta_{AICc}$ | *Weight* |
| --- | --- | --- | --- | --- |
| *Int + Emo* | 4 | 4339 | 0 | 0.59 |
| *Int + HRV + Emo* | 5 | 4341 | 2.243 | 0.19 |
| *Int + HRV + Emo + HRV \* Emo* | 6 | 4343 | 3.994 | 0.08 |
| *Int + HRV<sup>2</sup> + Emo + HRV<sup>2</sup> \* Emo* | 6 | 4343 | 3.994 | 0.08 |
| *Int + HRV + HRV<sup>2</sup> + Emo + HRV \* Emo + HRV<sup>2</sup> \* Emo* | 8 | 4344 | 5.03 | 0.04 |
| *Int* | 3 | 4463 | 123.8 | 0 |
| *Int + HRV<sup>2</sup>* | 4 | 4464 | 125.1 | 0 |
| *Int + HRV \* Emo* | 4 | 4464 | 125.3 | 0 |
| *Int + HRV* | 4 | 4465 | 126 | 0 |
| *Int + HRV + HRV<sup>2</sup>* | 5 | 4466 | 127.4 | 0 |

Note. *K* is the number of parameters in the model. *Int* = Intercept, *HRV* = resting HF-HRV, *Emo* = Type of emotion (present in Bal et al. (2010) versus not). All models include participants and items as random factors.

We then compared the parsimony of models containing main effects (HF-HRV and emotion type) and interaction effects. Model comparison showed no evidence in favor of a main effect of HF-HRV or toward an interaction between HRV and emotion compared to the intercept model, either for response times or accuracy (tables 3 and 4. HF-HRV did not predict performance in emotion identification. This absence of effect was observed regardless of the emotion type (“complex” versus “basic”). There was minimal evidence (*ER*_*i*_ = 0.631/0.211 =2.99 for response times and *ER*_*i*_ = 0.596/0.194 = 3.07 for accuracy) toward and principal effect of emotion type compared to the second best model and decisive evidence compared to the intercept model (Figure 5) with a marginal *R*^2^ of .06 and .05 respectively. Overall emotions absent from Bal et al. (2010) (i.e. “complex emotions”) were more difficult to identify compared to the emotion they used (i.e. “basic” emotions).

**Figure 5.**
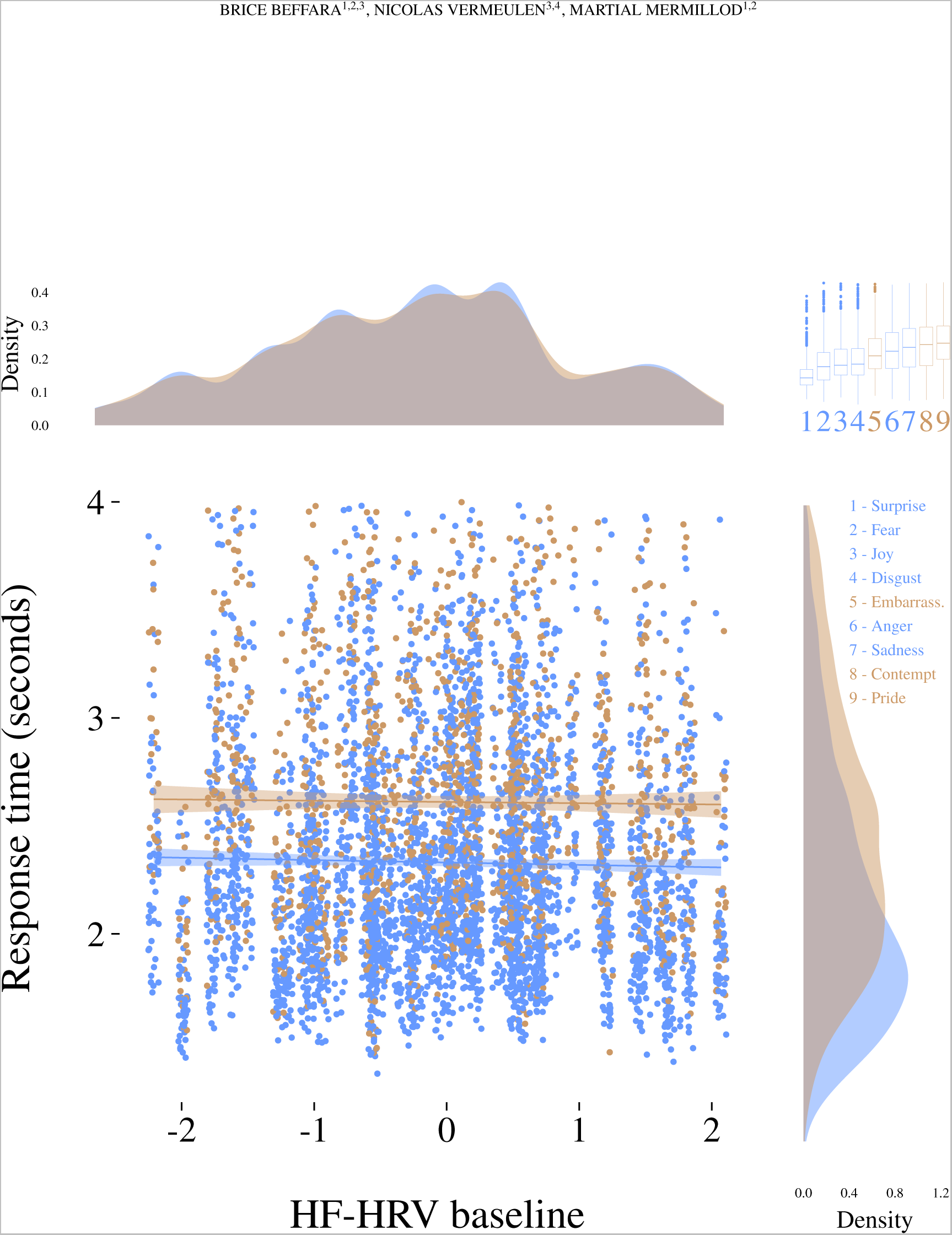
Response time for emotiosn identification as a function of resting HF-HRV and emotion type. Confidence regions represent 95% CIs. The top-right plot represents the ranking of the median response times relative to each EFEs.

## Discussion

We carried out a study in order to test whether HF-HRV was associated with better decoding of emotional facial expressions. Our protocol was built in order to combine the properties of previous studies on this subject (Bal et al., 2010; Quintana et al., 2012). We were able to measure reaction times and accuracy in an EFEs recognition task with both “basic” and “self-conscious” emotions (Schalk et al., 2011). In line with the observations of Bal et al. (2010), our results show that HF-HRV is not associated with better recognition of “basic”" emotions. While “self-conscious” emotions were harder to identify than “basic” emotions, the performance of participants was not predicted by HF-HRV. HF-HRV does not predict emotion identification on dynamic videos of whole faces, even taking difficulty into account.

The polyvagal theory predicts that the myelinated vagal connection between the heart and the brain can foster the perception of social cues in mammals (Porges, 2007). Quintana et al. (2012) showed that this feature of heart-brain interactions (as indexed by HF-HRV) is indeed associated with better performances at the RMET in healthy human adults. It is generally admitted that the RMET measures the ability to read others’ mental states. The association between HF-HRV and RMET performances can be interpreted as better emotion recognition skills in higher HF-HRV participants (Quintana et al., 2012). However, emotion recognition is not the only mechanism necessary to read other’s mental states. Attentional shifting and inhibition play a large part in Theory of Mind (ToM),i.e. the ability to attribute mental states to others (R. L. C. Mitchell & Phillips, 2015; Poletti, Enrici, & Adenzato, 2012; Samson, 2009). Several theoretical perspective have proposed a framework describing the interplay between emotion perception and ToM. Many of them propose the distinction between decoding the emotion from external stimulation and understanding its meaning for the other person (R. L. C. Mitchell & Phillips, 2015). This second step is likely to require inhibition of one’s perspective, rapid information updating, working memory, attentional switching between one’s and the other’s state (Carlson, Moses, & Breton, 2002; R. L. C. Mitchell & Phillips, 2015; Poletti et al., 2012; Samson, 2009).

As a consequence, the association between HF-HRV and mind reading could also be explained by better executive skills and not necessarily by better emotion identification abilities. Indeed, Bal et al. (2010) showed that HF-HRV was not associated with emotion recognition in healthy human children. They used an emotion categorization task with dynamic EFEs on six emotions (Porges et al., 2007). Still, it was not possible to put the work of Bal et al. (2010) and Quintana et al. (2012) in perspective because i) the population of interest was different (children vs. adults) and ii) the association between HF-HRV and RMET performances was observed when taking the items’ difficulty into account: it could be argued that the difficulty of the task in Bal et al. (2010) did not allow to discriminate the association with HF-HRV. We designed a study inspired from Bal et al. (2010) but tested healthy human adults and increased the difficulty of the task by adding three more EFEs to categorize. Model comparison by AICc showed that models without HF-HRV as a parameter were always far more parsimonious than models including HF-HRV as a parameter. This was observable for reaction times, accuracy, linear and quadratic shapes, even taking the difficulty of the task into account. This design allowed to discriminate between models with and without HF-HRV, that is to say, the parsimony of the models without HF-HRV was always clearly superior to the models with HF-HRV. This support the fact that HF-HRV is not associated with emotion recognition skills.

On the basis of these results, we propose that the association between HF-HRV and performances in “mental states” reading (Quintana et al., 2012) cannot be explained by better emotion recognition skills. The more plausible explanation at this stage would rather take attentional, working memory and executive skills into account. Interestingly, recent studies clearly show that higher HF-HRV individuals perform better in many cognitive tasks depending on executive and attentional functioning. The neurovisceral integration model provides a theoretical framework (Thayer & Lane, 2000) allowing to understand the association between HF-HRV and attention. The neural control of the heart is highly dependent on cortical inputs especially from the prefrontal cortex (PFC), the insula, and the anterior cingulate cortex (ACC). Variability observed in heart rate and mediated by the functioning of the myelinated vagus nerve is therefore largely influenced by attentional shifts, conflict monitoring, and inhibition. Conversely, it is also likely that afferent feed-backs from the heart can influence the central nervous system, therefore creating dynamic loops between the heart and the brain, fostering the adaptation of the organisms to internal and external demands. Neuroimaging studies bring evidence toward an important overlap between central nervous system activities associated with HRV (Thayer, Åhs, Fredrikson, Sollers, & Wager, 2012) and with ToM (Schurz, Radua, Aichhorn, Richlan, & Perner, 2014). The medial PFC (mPFC), the insula and the ACC play a large part in cardiovascular control and ToM. These areas show connections with the temporo-parietal junction (TPJ). It has been suggested that the TPJ is mainly involved in inferences about short-term intentions while more durable mental states could rather be taken over by the mPFC (Van Overwalle, 2009). The mPFC is also involved in inhibitory functions and interconnected with the ACC associated with cognitive control and conflict monitoring and with the insula underlying body states integration (Lane et al., 2009; Mier et al., 2010; Reeck, Ames, & Ochsner, 2016; Thayer & Lane, 2009). Therefore, brain areas involved in cardiovascular control and characterizing differences in HRV are often found associated with executive functioning, attentional regulation,and switching between one’s and other’s body states rather than emotion identification.

Even if we did not measure sensorimotor activity of the face during the tasks, we made the hypothesis that sensorimotor simulation would play an important part in the detection of emotions (Wood et al., 2016). This hypothesis was important in order to test the polyvagal proposition (Porges, 2001) according to which neural cardiovascular control is associated with neural sensorimotor control of the head and face muscles, both at an anatomical and at a functional level. In this perspective, our result does not validate that HF-HRV and sensorimotor skills are associated in order to perform a perceptive task such as decoding EFEs. Thus, it is plausible that HF-HRV predicts social skills (Beffara, Bret, Vermeulen, & Mermillod, 2016; Miller, Kahle, & Hastings, 2015) at another level. Attentional skills have already been suggested as the cognitive mechanism linking HF-HRV and social functioning (Keltner, Kogan, Piff, & Saturn, 2014). Obviously, we did not test this hypothesis in this study. However, as attention is a strong necessity to apply theory of mind (Lin, Keysar, & Epley, 2010) aside from decoding facial patterns, it is likely that the ability of higher HF-HRV individuals to process social signals is not due to better sensori-motor control but rather to better attentional or executive skills (Park & Thayer, 2014). Obviously, this proposition still needs to be specifically tested.

A solid set of studies highlight the association between HFHRV and working memory (Hansen, Johnsen, & Thayer, 2003; Hansen, Johnsen, Sollers, Stenvik, & Thayer, 2004), inhibition and attention switching (Kimhy et al., 2013), and more flexible attentional engagement and disengagement toward negative emotional stimuli (Park & Thayer, 2014; Park, Van Bavel, Vasey, Egan, & Thayer, 2012; Park, Vasey, Van Bavel, & Thayer, 2013). Consequently, whether at neuroimaging or behavioral level, better cognitive skills associated with higher resting state HF-HRV appear to be a more reliable candidate for explaining more accurate mind reading, while emotion identification abilities did not show substantial association with HF-HRV in our study. While further studies are needed to clearly establish the mediation of the HF-HRV – ToM link by executive functioning, we suggest that domain-general cognitive mechanisms (C. Heyes, 2014; Cecilia Heyes, 2016a, 2016b; Cecilia Heyes & Pearce, 2015) should be considered when studying in the functional association between HF-HRV and the social life.

### Conclusions

Heart-brain interactions are proposed to underlie socio-emotional skills (Porges, 2007). It has been shown that resting HF-HRV is associated with mental states reading (Quintana et al., 2012). These authors suggested that HF-HRV was linked to emotion recognition abilities. However, the current study does not allow to conclude that resting HF-HRV predict emotion recognition, even taking emotion type into account. Further studies should examine the role of executive functioning as a mediator of the HF-HRV – ToM association. Domain-general cognitive skills could account for the role of HF-HRV in social functioning.

## Aknowledgements

We thank Amélie Baldini and Elie Bes for technical support in data collection. We also thank all the participants for taking part in the study. We thank Pierre Maurage and Delphine Grynberg for their useful comments and constructive remarks. This research was funded by the french CNRS and the Institut Universitaire de France.

